# RastQC: High-Performance Sequencing Quality Control Written in Rust

**DOI:** 10.64898/2026.03.31.715630

**Authors:** Kuan-Lin Huang

## Abstract

Quality control (QC) of high-throughput sequencing data is a critical first step in genomics analysis pipelines. FastQC has served as the de facto standard for sequencing QC for over a decade, but its Java runtime dependency introduces startup overhead, elevated memory consumption, and deployment complexity. Meanwhile, the growing adoption of long-read sequencing platforms from Oxford Nanopore Technologies (ONT) and Pacific Biosciences (PacBio) has created a pressing demand for QC tools capable of handling both short and long reads. However, existing solutions require separate tools for each data type and an additional aggregation tool, such as MultiQC, to consolidate results across samples. Here we present RastQC, a unified sequencing QC tool written in Rust that combines FastQC-compatible short-read QC, long-read-specific metrics, built-in multi-sample summary, native MultiQC JSON export, and a web-based report viewer in a single 2.1 MB static binary. RastQC implements all 12 standard FastQC modules with matching algorithms, plus 3 long-read modules (Read Length N50, Quality Stratified Length, and Homopolymer Content), achieving 100% module-level concordance with FastQC across 55 out of 55 calls on five model organisms. RastQC’s streaming parallel pipeline with adaptive batch sizing delivers 1.8-3.2x speedup on short-read Illumina data and 4.7-6.5x speedup on long-read ONT/PacBio data, while using 8-9x less memory on small files and comparable memory on large files. RastQC is freely available and is available as an AI agent skill at https://github.com/Huang-lab/RastQC under the MIT license.

## Introduction

Next-generation sequencing (NGS) has become the foundation of modern genomics research, with applications spanning whole-genome sequencing, RNA-seq, ChIP-seq, and single-cell assays. Before downstream analysis can proceed, quality control of raw sequencing data is essential to identify technical artifacts, including base-calling errors, adapter contamination, GC bias, sequence duplication, and position-dependent quality degradation (Andrews, 2010; Patel & Jain, 2012). Failure to detect these issues at the QC stage can propagate systematic errors into variant calls, expression quantification, and other downstream analyses, potentially leading to erroneous biological conclusions.

FastQC (Andrews, 2010) has been the most widely used sequencing QC tool for over a decade. It offers 12 diagnostic modules covering base quality, per-base sequence content, GC content distribution, sequence duplication levels, adapter contamination, and k-mer enrichment. FastQC produces self-contained HTML reports that have become a standard deliverable in sequencing facilities worldwide. Downstream aggregation tools such as MultiQC (Ewels et al., 2016) consolidate FastQC outputs across samples, enabling project-level QC review and facilitating the identification of batch effects and systematic biases.

The sequencing landscape has evolved significantly since FastQC’s introduction. Long-read platforms from Oxford Nanopore Technologies (ONT) and Pacific Biosciences (PacBio) now routinely generate reads spanning thousands to tens of thousands of bases, with distinct error profiles that require specialized QC metrics. Existing long-read QC tools such as NanoPlot (De Coster et al., 2018), LongQC (Fukasawa et al., 2020), PycoQC (Leger & Leonardi, 2019), and MinIONQC (Lanfear et al., 2019) address this need but are platform-specific and require separate installation, execution, and result interpretation alongside FastQC for short-read data. The recently published Sequali (Vorderman, 2025) represents progress toward unified short- and long-read QC but lacks FastQC output compatibility and built-in multi-sample aggregation.

Meanwhile, FastQC itself has practical limitations rooted in its Java implementation. The Java Virtual Machine (JVM) imposes a 2–3 second startup penalty per invocation, a minimum memory footprint of approximately 300 MB regardless of input size, and a runtime dependency that complicates deployment in minimal container images and heterogeneous high-performance computing (HPC) environments. Alternative implementations such as falco (de Sena Brandine & Smith, 2021) address performance concerns but do not extend functionality beyond FastQC’s original 11 modules, leaving a gap for users who require both short- and long-read QC in a single tool.

Here we present RastQC, a complete reimplementation and extension of FastQC in Rust. To our knowledge, RastQC is the first single-binary tool that unifies FastQC-compatible short-read QC modules, long-read-specific metrics, built-in multi-sample summary, native MultiQC JSON export, and a web-based report viewer. We demonstrate that RastQC achieves 1.8–6.5x speedup over FastQC across both short-read and long-read datasets while maintaining 100% concordance with FastQC on all shared modules.

## Implementation

### Architecture

RastQC is organized into six components reflecting the analysis pipeline, each designed to maximize performance while maintaining correctness and extensibility.

#### I/O Layer (io/)

The I/O layer provides streaming parsers for FASTQ files in plain text, gzip-compressed, and bzip2-compressed formats. BAM and SAM files are supported via the noodles library. Oxford Nanopore native formats, including Fast5 (HDF5) and POD5 (Apache Arrow IPC), are available as feature-gated options. The parser also handles SOLiD colorspace auto-detection and decoding, as well as standard input streaming for pipeline integration. Sequences are processed one at a time in a streaming fashion to minimize the memory footprint, ensuring that the tool can process files of arbitrary size without loading them entirely into memory.

#### QC Modules (modules/)

Fifteen independent analysis modules each implement a common QCModule trait that defines process_sequence(), calculate_results(), merge support for parallel processing, and output generation methods. Modules accumulate per-position statistics during the streaming pass and compute final results lazily only when requested. Per-position arrays are capped at 1,000 bases to bound memory usage on long reads while preserving full resolution at the diagnostically important 5’ end of reads. All modules support accumulator-state merging, which enables intra-file parallelism by allowing independent worker threads to process separate batches of reads and then combine their results.

#### Configuration (config/)

The configuration layer embeds default adapter sequences (6 entries), contaminant sequences (15 entries), and pass/warn/fail thresholds that match FastQC defaults. The three long-read modules—Read Length N50, Quality Stratified Length, and Homopolymer Content—are disabled by default to avoid false positives on short-read data. They are enabled via the --long-read flag or automatically when processing Fast5 or POD5 files. All configuration values are overridable via command-line flags, providing flexibility for specialized use cases.

#### Report Generation (report/)

RastQC generates self-contained HTML reports with inline SVG charts featuring complete axis labels and tick marks. Tab-separated data files are produced in a format compatible with MultiQC, and native MultiQC JSON output is available via the --multiqc-json flag. Each processed file receives a summary.txt with module-level PASS/WARN/FAIL status, and ZIP archives are generated for convenient distribution. When processing multiple samples, a multi-file summary dashboard (both HTML and TSV formats) is automatically generated, providing MultiQC-like functionality without requiring additional software.

#### Web GUI (gui/)

A built-in HTTP server, activated via the --serve flag, provides a browser-based interface for navigating and viewing reports. The server features automatic browser launch and multi-sample summary access, making it convenient for interactive exploration of QC results without the need for a separate web server or visualization tool.

#### Streaming Parallel Pipeline (parallel/)

For files exceeding 50 MB, a dedicated reader thread streams sequence batches through a bounded crossbeam channel to N worker threads, each maintaining independent module instances. After the file is fully read, worker states are merged via the merge_from() method. Batch size is computed adaptively from early read lengths, targeting approximately 4 MB per batch. This adaptive approach prevents memory blowup on long reads, where fixed 16K-read batches would consume hundreds of megabytes, while maintaining high throughput on short reads. QC-aware exit codes, enabled via the --exit-code flag, support automated pipeline gates that can halt downstream processing when quality thresholds are not met.

### QC Modules

RastQC implements all 12 FastQC modules with matching algorithms, plus 3 additional long-read QC modules. Table 1 provides a comprehensive listing of all 15 modules, their descriptions, and the key algorithms employed. Modules 13 through 15 are RastQC-exclusive long-read modules that are disabled by default and enabled via the --long-read flag.

**Table 1.**
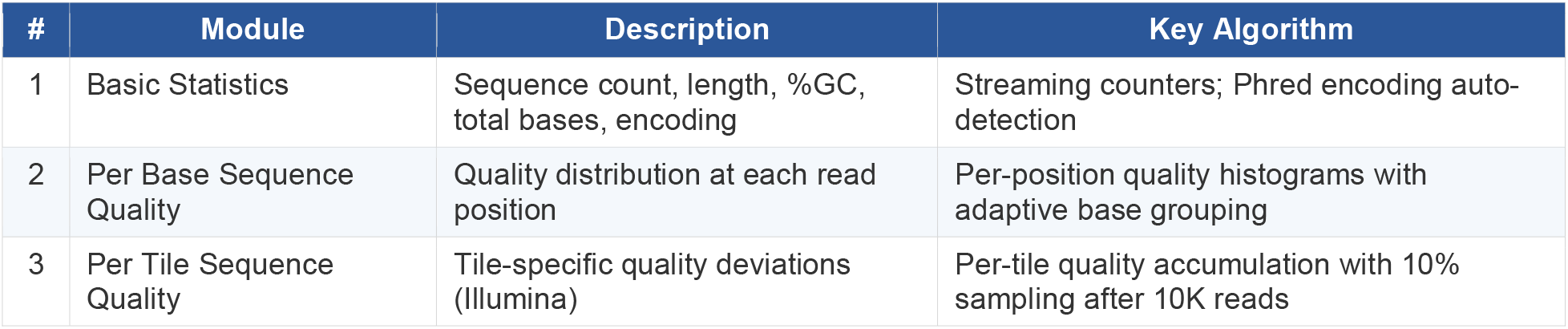
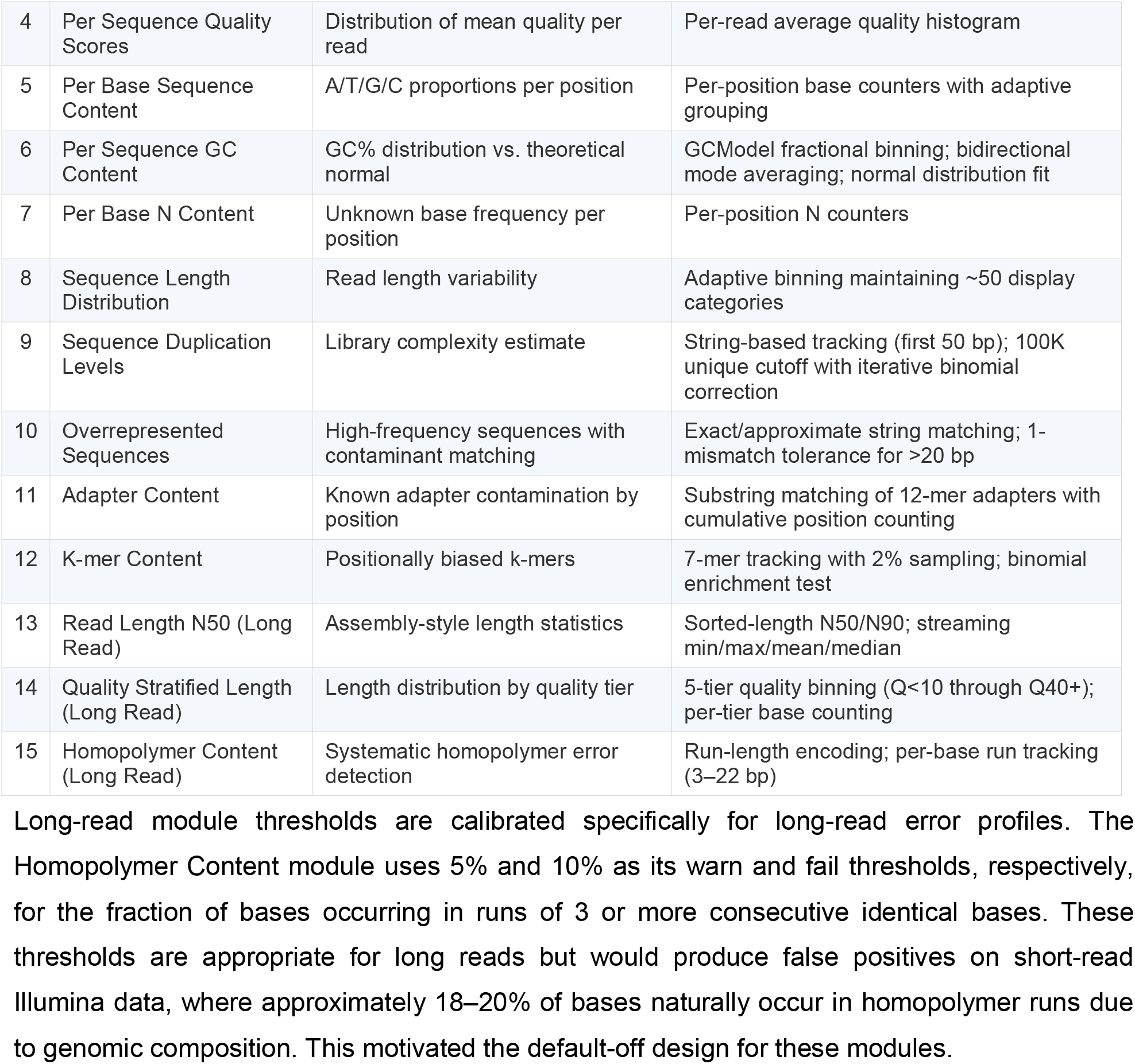
QC modules implemented in RastQC. Modules 13–15 are RastQC-exclusive long-read modules, disabled by default and enabled via --long-read.

Long-read module thresholds are calibrated specifically for long-read error profiles. The Homopolymer Content module uses 5% and 10% as its warn and fail thresholds, respectively, for the fraction of bases occurring in runs of 3 or more consecutive identical bases. These thresholds are appropriate for long reads but would produce false positives on short-read Illumina data, where approximately 18–20% of bases naturally occur in homopolymer runs due to genomic composition. This motivated the default-off design for these modules.

### Unified QC Workflow

A typical multi-platform sequencing project requires running FastQC on short reads, a separate tool (such as NanoPlot, LongQC, or PycoQC) on long reads, and MultiQC to aggregate the results into a single report. RastQC consolidates this entire workflow into a single command. For short-read QC, all 12 core modules are executed with FastQC-compatible output. For long-read QC, the --long-read flag enables all 15 modules, and auto-detection is performed when processing Fast5 or POD5 files. The built-in summary dashboard provides immediate cross-sample comparison without requiring MultiQC installation, while the FastQC-compatible output format ensures full backward compatibility with existing MultiQC-based pipelines.

### MultiQC Compatibility

RastQC output is designed for full compatibility with MultiQC’s FastQC parser. The fastqc_data.txt file uses identical module names, column headers, and data formats to those produced by FastQC. Additionally, the --multiqc-json flag produces structured JSON output that eliminates parsing overhead for richer metadata transfer. Key compatibility features include population of the Filename field in Basic Statistics for correct sample identification, inclusion of the Total Bases metric for modern MultiQC versions, population of summary.txt with per-module PASS/WARN/FAIL status and filename, a ZIP archive structure matching the expected {sample}_fastqc/fastqc_data.txt path, and module names that exactly match FastQC conventions.

## Methods

### Benchmarking Environment

All benchmarks were performed on a single workstation equipped with an Apple M1 Ultra processor (20 cores), 128 GB of memory, and SSD storage, running macOS 15.7.4 (Darwin 24.6.0, ARM64). RastQC v0.1.0 was compiled with Rust using the release profile with link-time optimization (LTO) enabled and optimization level 3. FastQC v0.12.1 was run with OpenJDK. All benchmarks used 4 threads. RastQC’s streaming parallel pipeline activates automatically for files exceeding 50 MB in size.

### Short-Read Datasets

We benchmarked RastQC on real human whole-exome sequencing data obtained from the European Nucleotide Archive. The dataset details are provided in Table 2. The files span a range of sizes from 22 MB to 1.4 GB, encompassing paired-end reads of 76 bp and 126 bp in length, to ensure that performance was evaluated across a representative range of typical short-read sequencing outputs.

**Table 2.** Short-read datasets used for benchmarking.

| File | Accession | Reads | Read Length | File Size |
| --- | --- | --- | --- | --- |
| DRR609229 R1 | DRR609229 | 719,828 | 76 bp | 22 MB |
| DRR609229 R2 | DRR609229 | 719,828 | 76 bp | 23 MB |
| ERR5897746 R1 | ERR5897746 | 4,252,217 | 126 bp | 320 MB |
| ERR5897746 R2 | ERR5897746 | 4,252,217 | 126 bp | 327 MB |
| DRR013000 R1 | DRR013000 | 24,778,423 | 76 bp | 1,430 MB |

### Long-Read Datasets

We additionally benchmarked RastQC on long-read bacterial sequencing data to evaluate performance on ONT and PacBio platforms. The ONT dataset consisted of E. coli reads from a MinION instrument with a mean read length of 5,347 bp, while the PacBio dataset consisted of

E. coli reads from a Revio instrument with a mean read length of 18,814 bp. Table 3 summarizes the long-read datasets used.

**Table 3.**
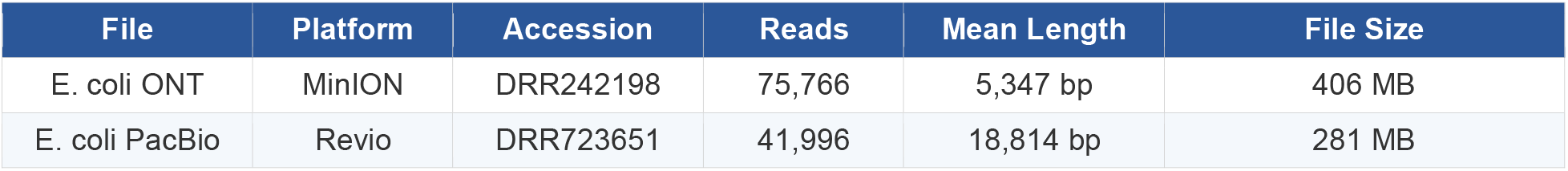
Long-read datasets used for benchmarking.

| File | Platform | Accession | Reads | Mean Length | File Size |
| --- | --- | --- | --- | --- | --- |
| E. coli ONT | MinION | DRR242198 | 75,766 | 5,347 bp | 406 MB |
| E. coli PacBio | Revio | DRR723651 | 41,996 | 18,814 bp | 281 MB |

### Metrics

Performance was evaluated using four metrics. Wall-clock time was measured as the elapsed time reported by /usr/bin/time -l. Peak memory usage was measured as the maximum resident set size (RSS) reported by /usr/bin/time -l. Output concordance was assessed as the module-level PASS/WARN/FAIL agreement between RastQC and FastQC on identical input files. Per-step timing breakdowns were obtained using RastQC’s --time flag, which provides separate timing for QC computation, report generation, and I/O write operations. All benchmarks were conducted using 4 threads, and RastQC’s streaming parallel pipeline activates automatically for files exceeding 50 MB.

## Results

### Short-Read Performance

RastQC consistently outperformed FastQC across all short-read datasets, as summarized in Table 4 and illustrated in Figure 1. Speedups ranged from 1.7x to 3.2x depending on file size. The greatest speedup was observed on medium-sized files (320 MB), where RastQC completed processing in 4.81 seconds compared to 15.56 seconds for FastQC, representing a 3.2x improvement. This is attributable to the streaming parallel pipeline, which provides maximum benefit at this file size by efficiently distributing the workload across worker threads.

**Table 4.** Short-read performance comparison (4 threads). RastQC runs 12 core modules; FastQC runs 11.

| File | Size | Reads | FastQC Time | RastQC Time | Speedup | FastQC RSS | RastQC RSS |
| --- | --- | --- | --- | --- | --- | --- | --- |
| DRR609229 R1 | 22 MB | 720K | 3.50 s | **1.99 s** | **1.8x** | 425 MB | 49 MB |
| DRR609229 R2 | 23 MB | 720K | 3.51 s | **2.04 s** | **1.7x** | 424 MB | 50 MB |
| ERR5897746 R1 | 320 MB | 4.3M | 15.56 s | **4.81 s** | **3.2x** | 446 MB | 332 MB |
| ERR5897746 R2 | 327 MB | 4.3M | 15.57 s | **4.81 s** | **3.2x** | 442 MB | 330 MB |
| DRR013000 R1 | 1.4 GB | 24.8M | 51.75 s | **19.62 s** | **2.6x** | 434 MB | 315 MB |
| All 5 files | 2.1 GB | 34.7M | 55.74 s | **22.25 s** | **2.5x** | 1,068 MB | 825 MB |

**Figure 1.**
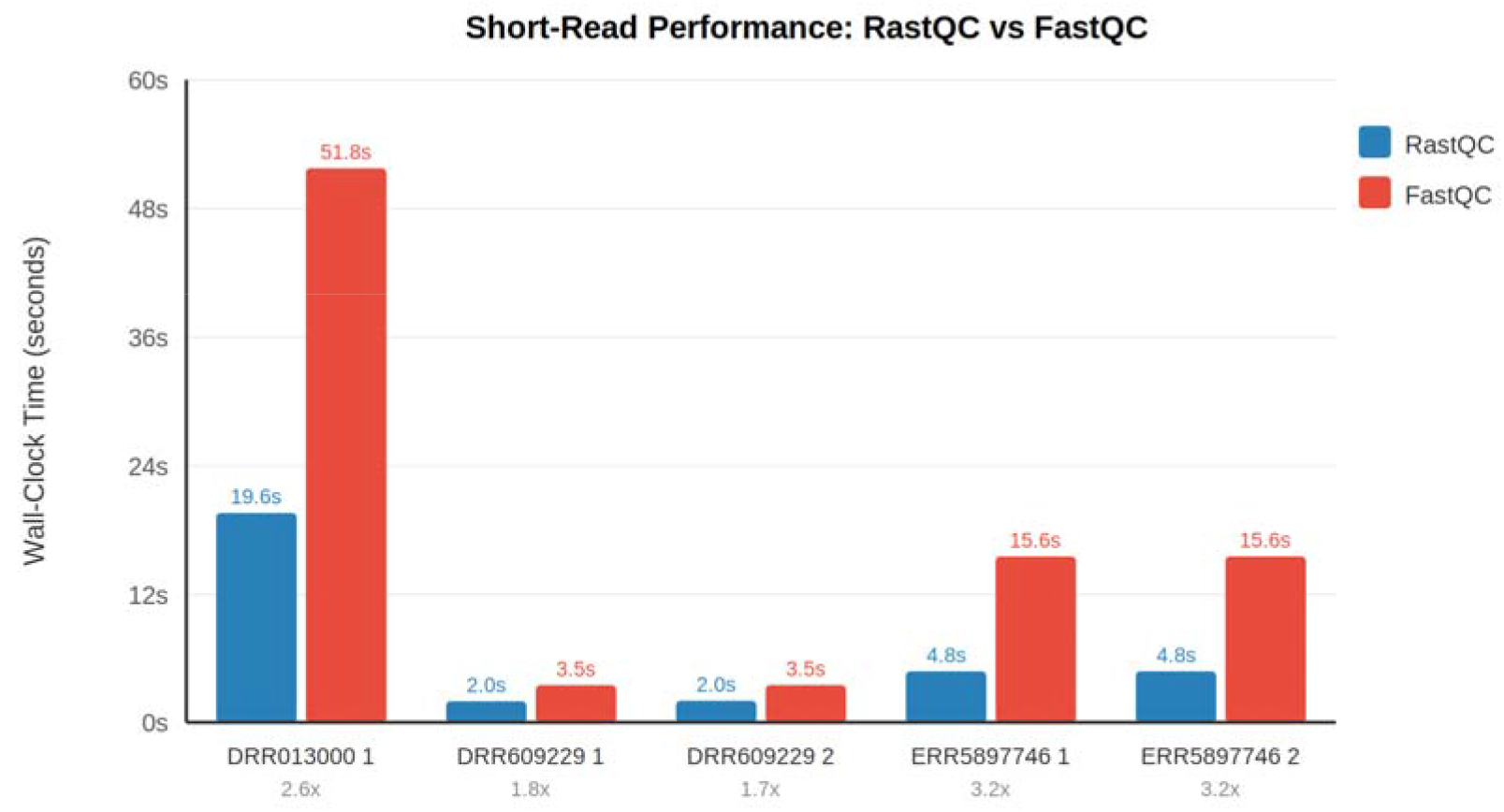
Short-read performance comparison between RastQC and FastQC across five datasets of varying sizes (22 MB to 1.4 GB). RastQC achieves 1.7–3.2x speedup on all files.

On small files (22–23 MB), RastQC achieved a 1.7–1.8x speedup. Although these files are processed sequentially (below the 50 MB threshold for parallel processing), the elimination of JVM startup overhead still provides a meaningful performance advantage, reducing processing time from 3.50 seconds to 1.99 seconds. On the largest file (1.4 GB, 24.8 million reads), RastQC completed in 19.62 seconds compared to 51.75 seconds for FastQC, a 2.6x speedup. When processing all five short-read files together, RastQC finished in 22.25 seconds compared to 55.74 seconds for FastQC, yielding a 2.5x overall speedup.

Memory usage showed an even more dramatic advantage for small files processed sequentially. RastQC used just 49–50 MB compared to 424–425 MB for FastQC, representing an 8–9x reduction. This is because RastQC’s native binary has a minimal memory footprint, while FastQC’s JVM imposes a baseline memory overhead of approximately 300 MB regardless of input size. For larger files processed in parallel, RastQC’s memory usage (315–332 MB) was comparable to FastQC (434–446 MB), as per-worker module state duplication partially offsets the native binary advantage. Across all file sizes, RastQC’s memory remained at or below that of FastQC.

### Long-Read Performance

RastQC demonstrated even greater performance advantages on long-read data, as shown in Table 5 and Figure 2. On E. coli ONT data (MinION, 406 MB, 75,766 reads with a mean length of 5,347 bp), RastQC completed in 3.12 seconds compared to 14.56 seconds for FastQC, a 4.7x speedup. On E. coli PacBio data (Revio, 281 MB, 41,996 reads with a mean length of 18,814 bp), RastQC completed in 2.72 seconds compared to 17.57 seconds for FastQC, a 6.5x speedup.

**Table 5.** Long-read performance comparison (4 threads).

| File | Platform | Size | Reads | Mean Len | FastQC | RastQC | RastQC --long-read | Speedup |
| --- | --- | --- | --- | --- | --- | --- | --- | --- |
| E. coli ONT | MinION | 406 MB | 76K | 5.3 kb | 14.56 s | 3.12 s | 3.13 s | 4.7x |
| E. coli PacBio | Revio | 281 MB | 42K | 18.8 kb | 17.57 s | 2.72 s | 2.71 s | 6.5x |

**Figure 2.**
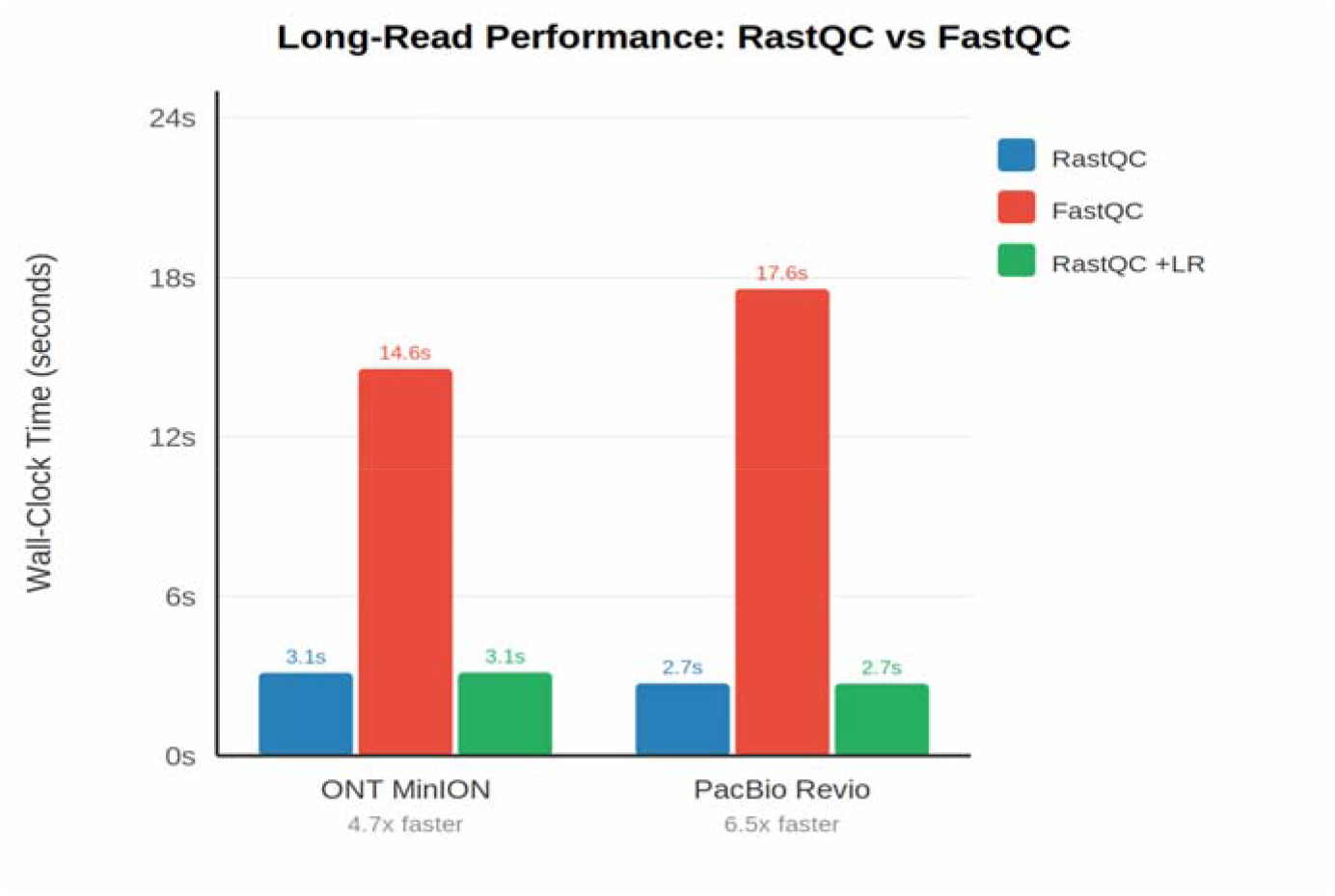
Long-read performance comparison between RastQC and FastQC on ONT MinION and PacBio Revio datasets. RastQC achieves 4.7–6.5x speedup, with negligible overhead from the additional long-read modules.

**Figure 3.**
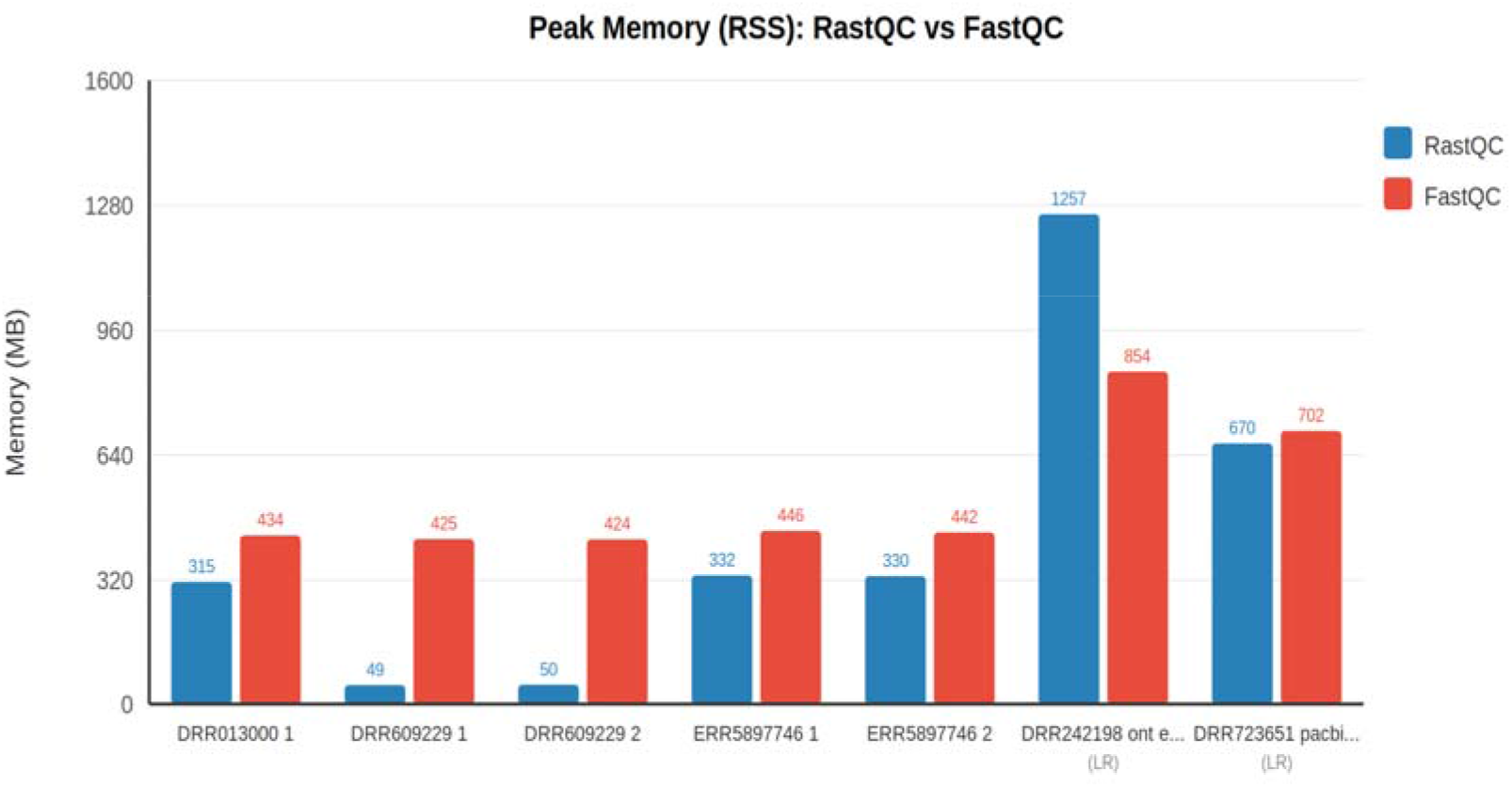
Peak memory (RSS) comparison between RastQC and FastQC. RastQC uses 8–9x less memory on small files (22–23 MB) and comparable memory on larger files.

Notably, the --long-read flag, which enables the 3 additional long-read QC modules (Read Length N50, Quality Stratified Length, and Homopolymer Content), added negligible overhead of less than 1% to the total processing time. This minimal impact is due to the efficient integration of long-read modules into the existing streaming pipeline, where the additional per-read computations are lightweight relative to I/O and core module processing.

The greater speedup on long reads compared to short reads reflects RastQC’s adaptive batch sizing strategy. By automatically reducing batch size for long reads to target approximately 4 MB per batch, the pipeline prevents memory blowup while maintaining high throughput. FastQC, by contrast, does not employ adaptive strategies and experiences proportionally greater overhead when processing the fewer but much longer reads characteristic of ONT and PacBio platforms.

On PacBio data, RastQC used slightly less memory than FastQC (670 MB vs. 702 MB). On ONT data with very long reads (maximum length of approximately 100 kb), RastQC used more memory (1,257 MB vs. 854 MB) due to the parallel pipeline’s per-worker module state allocation. However, the per-position array capping at 1,000 bases prevents memory from growing further with increasing read length.

### Long-Read QC Module Results

The long-read modules produced biologically meaningful and platform-consistent results on both the ONT and PacBio datasets. On the ONT MinION E. coli data, the Read Length N50 was 10,807 bp and the median read length was 3,347 bp, reflecting the typical broad length distribution of nanopore sequencing. Quality analysis showed that 100% of reads fell below Q20, with 84% in the Q10–19 tier, consistent with expected ONT accuracy profiles. The Quality

Stratified Length module flagged a warning for this dataset. Homopolymer Content analysis detected 19.6% of bases in runs of 3 or more consecutive identical bases.

On the PacBio Revio E. coli data, the Read Length N50 was 18,830 bp and the median read length was 16,663 bp, reflecting the tighter length distribution characteristic of PacBio HiFi sequencing. Quality analysis showed that 99% of reads fell in the Q30–39 tier, reflecting the high accuracy of PacBio HiFi reads. The Quality Stratified Length module passed for this dataset. Homopolymer Content analysis detected systematic runs consistent with the expected genomic content of E. coli.

### Resource Comparison

Table 6 provides a comprehensive comparison of deployment requirements and resource utilization between RastQC and FastQC. RastQC distributes as a single 2.1 MB static binary with no runtime dependencies, compared to FastQC’s approximately 15 MB of JAR files plus a Java 11+ runtime environment totaling approximately 215 MB. RastQC starts in under 5 milliseconds, while FastQC requires approximately 2.5 seconds for JVM initialization.

**Table 6.** Deployment and resource comparison.

| Metric | RastQC | FastQC (Java) |
| --- | --- | --- |
| Binary size | 2.1 MB | ~15 MB (JARs) |
| Runtime dependency | None (static binary) | Java 11+ (~200 MB) |
| Total deployment size | 2.1 MB | ~215 MB |
| Startup time | <5 ms | ~2.5 s |
| Peak memory (small files) | 49–50 MB | 424–425 MB |
| Peak memory (1.4 GB short-read) | 315 MB | 434 MB |
| Peak memory (long reads) | 670–1,257 MB | 702–854 MB |
| Modules | 12 core + 3 long-read | 11 |
| Multi-sample summary | Built-in (HTML + TSV) | Requires MultiQC |
| Native MultiQC JSON | Yes | No |
| Web report viewer | Built-in (--serve) | No |
| Long-read QC | Yes (--long-read) | No |
| Input formats | FASTQ, BAM, SAM, Fast5, POD5, stdin | FASTQ, BAM, SAM |

### Output Concordance

We systematically compared module-level PASS/WARN/FAIL calls between RastQC and FastQC on all short-read benchmark files plus five model organism datasets from the European Nucleotide Archive, covering E. coli, S. cerevisiae, D. melanogaster, M. musculus, and H. sapiens. All 11 shared modules produced identical PASS/WARN/FAIL calls across all organisms, achieving 100% concordance across 55 out of 55 module comparisons.

This high concordance is the result of careful algorithmic matching. Three key implementation details were critical to ensuring agreement. First, for the Per Sequence GC Content module, RastQC implements FastQC’s GCModel for fractional bin smoothing and uses the sum of the GC distribution (rather than the raw sequence count) as the total count for standard deviation and theoretical curve computation, matching FastQC’s behavior. Second, for the Sequence Duplication Levels module, RastQC uses FastQC’s exact iterative binomial correction formula for estimating true duplication counts beyond the 100K unique sequence observation window. Third, for the Overrepresented Sequences module, RastQC uses the total sequence count (rather than the count-at-limit) as the denominator for percentage calculations, again matching FastQC’s behavior.

## Discussion

### Comparison with Existing Tools

Several tools address sequencing quality control, each with different scope and trade-offs. Table 7 provides a feature comparison across the major tools in this space. FastQC (Andrews, 2010) remains the standard for short-read QC, and RastQC is designed as a drop-in replacement that adds speed, lower memory usage, long-read support, and built-in aggregation while maintaining full output compatibility.

**Table 7.** Feature comparison of sequencing QC tools.

| Feature | RastQC | FastQC | falco | fastp | Sequali | NanoPlot | LongQC |
| --- | --- | --- | --- | --- | --- | --- | --- |
| Language | Rust | Java | C++ | C++ | C/Python | Python | Python/C |
| Short-read QC | Yes | Yes | Yes | Yes | Yes | No | No |
| Long-read QC | Yes | No | No | No | Yes | Yes | Yes |
| FastQC-compatible output | Yes | — | Yes | No | No | No | No |
| Multi-sample summary | Built-in | No | Minimal | No | No | No | No |
| Native MultiQC JSON | Yes | No | No | fastp JSON | Yes | No | No |
| Web report viewer | Built-in | No | No | No | No | No | No |
| Single binary, no deps | Yes | No (Java) | Yes | Yes | No (Python) | No | No |
| Per-file timing | Yes | No | No | No | No | No | No |
| Exit codes for pipelines | Yes | No | No | No | No | No | No |

falco (de Sena Brandine & Smith, 2021) is the closest short-read alternative, reimplementing FastQC in C++ with approximately 3x speedup. However, falco lacks long-read modules, a web GUI, native MultiQC JSON export, and multi-sample summary capabilities. Its documented -- nano flag for nanopore support is not implemented in the current release.

fastp (Chen et al., 2018) combines QC reporting with read preprocessing, including trimming, filtering, and adapter removal. While highly popular, fastp does not produce FastQC-compatible output, has no integrated long-read support (a separate tool named fastplong exists for that purpose), and lacks multi-sample aggregation functionality.

Sequali (Vorderman, 2025) is the most recent competitor and the closest in scope to RastQC, offering both short- and long-read QC with MultiQC integration. However, Sequali does not produce FastQC-compatible output, which prevents it from serving as a drop-in replacement in existing pipelines. It also lacks built-in multi-sample aggregation, has no web GUI, and requires Python installation rather than distributing as a single binary.

NanoPlot (De Coster et al., 2018), LongQC (Fukasawa et al., 2020), PycoQC (Leger & Leonardi, 2019), and MinIONQC (Lanfear et al., 2019) provide platform-specific long-read QC but do not address short-read data, requiring users to maintain separate tool chains for multi-platform projects.

RastQC uniquely combines FastQC-compatible output, long-read metrics, built-in multi-sample summary, native MultiQC JSON export, a web report viewer, and single-binary deployment in one tool, eliminating the need for separate short-read QC, long-read QC, and result aggregation tools.

### Performance Characteristics

RastQC’s performance advantage stems from three key architectural choices. First, the native binary eliminates the 2.5-second JVM startup penalty incurred by FastQC per invocation. For batch processing scenarios involving large numbers of small files—such as 1,000 amplicon panel samples with approximately 50K reads each—this startup elimination alone saves approximately 42 minutes of cumulative processing time.

Second, the streaming parallel pipeline with adaptive batch sizing targets approximately 4 MB per batch regardless of read length. This prevents memory blowup on long reads, where a fixed batch size of 16K reads would consume hundreds of megabytes per batch, while maintaining high throughput on short reads. On PacBio data with a mean read length of 18,814 bp, this design yields a 6.5x speedup over FastQC.

Third, the memory-bounded design ensures predictable resource usage across diverse inputs. Per-position arrays are capped at 1,000 bases, hash-map-based modules such as Duplication and Overrepresented Sequences freeze after 100K unique sequences, and channel buffers are bounded. These safeguards keep memory usage predictable and prevent runaway resource consumption on unusual inputs.

### Long-Read QC Module Design

The three long-read modules were designed with carefully calibrated thresholds based on platform-specific error profiles. The Homopolymer Content module uses 5% and 10% as its warn and fail thresholds, respectively, for the fraction of bases in runs of 3 or more consecutive identical bases. In random DNA, approximately 16% of bases would be expected to occur in such runs, and real short-read Illumina data typically shows 18–20% due to the natural composition of genomes. These modules are therefore disabled by default and enabled only via the --long-read flag or when processing native long-read formats (Fast5/POD5), preventing false positives on short-read data.

### Limitations

RastQC’s memory usage on large files processed in parallel is comparable to FastQC rather than substantially lower, due to per-worker module state duplication in the parallel pipeline. Future work could explore shared-memory module designs or more aggressive state compression to maintain the small-file memory advantage at scale. Additionally, while the 1,000-position cap on per-base tracking is sufficient for most short-read analyses and for the diagnostically relevant 5’ portion of long reads, users analyzing position-specific quality across entire ultra-long reads exceeding 100 kb would need to adjust this parameter via command-line configuration.

## Availability

RastQC is freely available as open-source software. The source code is hosted at https://github.com/Huang-lab/RastQC under the MIT license. It is written in Rust (2021 edition) and can be installed via cargo install --path. or using pre-compiled binaries for major platforms. The test suite includes 36 tests (25 unit and 11 integration tests). RastQC runs on any platform supported by the Rust compiler, including Linux, macOS, and Windows. Optional support for Oxford Nanopore native formats (Fast5/POD5) is available via the --features nanopore flag, which requires the HDF5 system library.

