## Supplementary Information for "RastQC: High-Performance Sequencing Quality Control Written in Rust"

Kuan-Lin Huang

### **S1. Detailed Algorithmic Specifications for QC Modules**

This section provides the precise algorithmic details for each of the 15 QC modules implemented in RastQC. These specifications document the exact procedures used to ensure 100% concordance with FastQC on all 11 shared modules and describe the novel long-read modules in full.

#### **S1.1 Basic Statistics (Module 1)**

The Basic Statistics module accumulates global counters during the single-pass streaming phase. For each read, the module increments the total sequence counter, adds the read length to the total base counter, and updates minimum and maximum observed read lengths. GC content is computed by counting G and C bases per read and accumulating a global sum; the final %GC is reported as the ratio of total G+C bases to total A+T+G+C bases, rounded to the nearest integer. Phred encoding is auto-detected from the first 10,000 quality scores: if any quality character has an ASCII value below 59 (corresponding to Phred+33 encoding), Sanger/Illumina 1.9+ encoding is assumed; otherwise, Illumina 1.5+ encoding (Phred+64) is reported. The filename field is populated from the input path basename, which is critical for correct sample identification in downstream MultiQC aggregation.

#### **S1.2 Per Base Sequence Quality (Module 2)**

This module maintains per-position quality histograms for Phred scores 0 through 93. For each read, the quality value at each position is recorded in a two-dimensional array indexed by position and quality score. To manage memory and maintain visual clarity, positions beyond a configurable threshold are grouped using FastQC’s adaptive base grouping scheme. The grouping algorithm divides positions into bins of increasing width: positions 1–10 are reported individually, positions 11–50 are grouped in bins of 5, positions 51–100 in bins of 10, positions 101–500 in bins of 50, and positions beyond 500 in bins of 100. For each group, the module reports the mean, median, lower quartile (Q1), upper quartile (Q3), and the 10th and 90th percentiles of the quality distribution.

The pass/warn/fail thresholds are as follows: a PASS is assigned if the lower quartile (Q1) at every position is 10 or above and the median at every position is 25 or above. A WARN is issued if the lower quartile at any position falls below 10 or the median at any position falls below 25. A FAIL is triggered if the lower quartile at any position falls below 5 or the median at any position falls below 20.

#### **S1.3 Per Tile Sequence Quality (Module 3)**

This module is specific to Illumina data where sequence identifiers contain tile information. For each tile, mean quality at each position is compared to the global mean quality at that position to compute a deviation score. Tile identifiers are extracted from the read header using pattern matching for Illumina-format headers (colon-delimited fields). To reduce computational overhead without sacrificing statistical power, this module employs a two-phase sampling strategy: all reads are processed for the first 10,000 sequences, after which a 10% random sampling rate is applied. The sampling is implemented using a deterministic hash of the read index modulo 10. This provides robust tile-level statistics even for very large files while bounding computational cost.

The pass/warn/fail thresholds compare tile-level deviations to the global mean. A WARN is issued if any tile at any position deviates by more than 2 Phred scores from the global mean. A FAIL is triggered if any tile at any position deviates by more than 5 Phred scores.

#### **S1.4 Per Sequence Quality Scores (Module 4)**

For each read, the mean quality score is computed as the arithmetic mean of all quality values across the read, rounded to the nearest integer. The module accumulates a histogram of these per-read mean quality values. The pass/warn/fail assessment is based on the mode of this distribution: a FAIL is assigned if the most commonly observed mean quality is below 20, a WARN if below 27, and a PASS otherwise.

#### **S1.5 Per Base Sequence Content (Module 5)**

This module tracks the proportion of A, T, G, and C bases at each read position. Per-position base counters are maintained and converted to percentages at reporting time. The same adaptive base grouping scheme used in Module 2 is applied here. The module evaluates whether the difference between A and T percentages or between G and C percentages exceeds a threshold at any position. A WARN is triggered if the difference exceeds 10 percentage points, and a FAIL is triggered if the difference exceeds 20 percentage points. These thresholds detect library preparation biases that produce non-random base composition at specific positions, such as hexamer priming bias commonly observed at the 5’ end of RNA-seq reads.

#### **S1.6 Per Sequence GC Content (Module 6)**

The GC content module computes the GC percentage for each read and builds a histogram of these values from 0% to 100%. RastQC implements FastQC’s GCModel class for fractional bin smoothing: when a read’s GC percentage falls between integer bins, the count is distributed fractionally between the two adjacent bins. This smoothing prevents quantization artifacts that would otherwise produce a jagged histogram, particularly for short reads where each base represents a large fraction of total content.

A theoretical normal distribution is fitted to the observed GC distribution. The mean is computed using bidirectional mode averaging: starting from the distribution’s mode, the algorithm walks left and right until the histogram value drops below 50% of the mode peak, averaging the positions at which this occurs. The standard deviation is computed from the GC distribution using the sum of the distribution (not the raw sequence count) as the denominator, matching FastQC’s behavior. This distinction is critical for concordance because fractional bin smoothing redistributes counts such that the sum of the GC distribution may differ slightly from the raw sequence count. The deviation from the theoretical curve is quantified as the sum of absolute differences, normalized by total count. A WARN is issued if this normalized deviation exceeds 15%, and a FAIL if it exceeds 30%.

#### **S1.7 Per Base N Content (Module 7)**

This module tracks the percentage of N (unknown/ambiguous) bases at each read position. Per-position N counters are maintained alongside total base counters, and the N percentage is computed at reporting time. The adaptive base grouping scheme is applied. A WARN is issued if the N percentage exceeds 5% at any position, and a FAIL is triggered if it exceeds 20%. Elevated N content typically indicates base-calling failures or sequencing chemistry problems at specific cycle positions.

#### **S1.8 Sequence Length Distribution (Module 8)**

The Sequence Length Distribution module records the frequency of each observed read length. For datasets with highly variable read lengths (common in long-read sequencing), an adaptive binning algorithm maintains approximately 50 display categories by merging adjacent bins. The module reports whether all reads have the same length (common for Illumina data) or whether there is length variability. A WARN is issued if not all sequences are the same length (which is expected for quality-trimmed data), and a FAIL is triggered if any sequences have zero length.

#### **S1.9 Sequence Duplication Levels (Module 9)**

The Sequence Duplication Levels module estimates library complexity by tracking how many times each unique sequence appears. Due to the computational infeasibility of tracking all unique sequences in large libraries, RastQC employs the same approach as FastQC: only the first 100,000 unique sequences are tracked in a hash map. For each incoming sequence, only the first 50 bp are used as the deduplication key (to normalize for 3’ quality trimming variability). Sequences are reverse-complemented and the lexicographically smaller of the forward and reverse-complement forms is used as the canonical key.

After the 100K unique sequence limit is reached, the hash map is frozen—no new unique sequences are added, but counts for existing sequences continue to be incremented. To correct for the sampling bias introduced by this observation window, FastQC’s iterative binomial correction formula is applied. The correction estimates the true number of unique sequences that would have been observed at each duplication level if all sequences had been tracked. Specifically, for a file with T total sequences and S observed unique sequences in the 100K window, the expected true count of sequences at duplication level d is estimated by solving the binomial expectation equation iteratively. RastQC implements this exact formula to ensure concordance.

The module reports two metrics: the percentage of sequences remaining after deduplication and the distribution of duplication levels (1 through 10+, and >500, >1K, >5K, >10K, >50K, >100K). A WARN is issued if non-unique sequences constitute more than 20% of the total, and a FAIL is triggered if they constitute more than 50%.

#### **S1.10 Overrepresented Sequences (Module 10)**

This module identifies sequences that appear at unusually high frequency, which may indicate adapter contamination, ribosomal RNA, or PCR artifacts. The implementation tracks exact sequence occurrences using the same 100K unique sequence hash map as the Duplication module (the first 50 bp of each read). Sequences that constitute more than 0.1% of total reads are reported as overrepresented.

Each overrepresented sequence is matched against a built-in contaminant database containing 15 common contaminant sequences (adapter dimers, primer sequences, ribosomal RNA fragments). Matching is performed using both exact substring matching and approximate matching with 1-mismatch tolerance for sequences longer than 20 bp. The percentage calculation uses the total sequence count as the denominator (not the count at the point the hash map was frozen), which is critical for concordance with FastQC. A WARN is issued if any single sequence exceeds 0.1% of total reads, and a FAIL if any exceeds 1%.

#### **S1.11 Adapter Content (Module 11)**

The Adapter Content module detects the presence of known sequencing adapter sequences at each read position. RastQC searches for 6 built-in adapter sequences by default, matching the FastQC adapter set: Illumina Universal Adapter, Illumina Small RNA 3’ Adapter, Illumina Small RNA 5’ Adapter, Nextera Transposase Sequence, SOLID Small RNA Adapter, and TruSeq Adapter Index 1. The detection algorithm performs substring matching using 12-mer subsequences of each adapter: for each position in a read, the algorithm checks whether any 12-mer adapter fragment begins at that position. When a match is found, all downstream positions (to the end of the read) are counted as adapter-contaminated, implementing cumulative position counting. A WARN is issued if any position shows more than 5% adapter content, and a FAIL if any position exceeds 10%.

#### **S1.12 K-mer Content (Module 12)**

The K-mer Content module identifies k-mers (short nucleotide subsequences) that are enriched at specific positions relative to their overall frequency. RastQC tracks all 7-mers (16,384 possible sequences) and their positional distribution. To manage computational cost, a 2% sampling rate is applied—only 1 in every 50 reads is analyzed for k-mer content. For each sampled read, all 7-mers are extracted and their positions recorded. After processing, the observed positional count for each k-mer is compared to the expected count (based on the k-mer’s overall frequency) using a binomial enrichment test. K-mers with statistically significant positional enrichment (p < 0.01 after Bonferroni correction) are reported. A WARN is issued if any k-mer shows a position-specific enrichment ratio of 3 or greater, and a FAIL if the ratio exceeds 5.

#### **S1.13 Read Length N50 (Module 13, Long-Read)**

The Read Length N50 module computes assembly-style length statistics that are standard metrics in the long-read sequencing community. During the streaming pass, the module records the length of every read in a vector. At reporting time, lengths are sorted in descending order and the cumulative sum is computed. N50 is defined as the read length at which 50% of the total base count is contained in reads of that length or longer. N90 is defined analogously at the 90% threshold. Additionally, the module reports minimum, maximum, mean, and median read lengths. These metrics provide immediate insight into the yield and quality of a long-read sequencing run. A WARN is issued if the N50 is less than 1,000 bp, and a FAIL if it is less than 500 bp.

#### **S1.14 Quality Stratified Length (Module 14, Long-Read)**

This module partitions reads into five quality tiers based on their mean Phred quality score and reports the total base count contributed by each tier. The five tiers are: Q < 10, Q10–19, Q20–29, Q30–39, and Q40+. For each read, the mean quality is computed and the read’s total bases are added to the appropriate tier’s counter. This analysis reveals whether a long-read dataset’s yield is concentrated in high-quality or low-quality reads, which has direct implications for downstream analysis sensitivity. For example, ONT data typically shows the majority of bases in the Q10–19 tier, while PacBio HiFi data concentrates bases in the Q30–39 tier. A WARN is issued if more than 50% of total bases fall in reads with mean quality below Q10, and a FAIL if more than 80% of bases are below Q10.

#### **S1.15 Homopolymer Content (Module 15, Long-Read)**

The Homopolymer Content module detects systematic homopolymer runs that may indicate sequencing errors characteristic of long-read platforms. The algorithm applies run-length encoding (RLE) to each read: consecutive identical bases are collapsed into a single run characterized by the base identity (A, T, G, or C) and run length. Runs of length 3 through 22 are tracked per base, producing a 4 × 20 count matrix. The primary metric is the fraction of total bases that occur in homopolymer runs of 3 or more bases.

In random DNA with equal base frequencies, the expected fraction of bases in homopolymer runs of 3+ is approximately 16.0%. In real genomic DNA, this fraction varies by organism and is typically 16–22% for short-read Illumina data. Long-read platforms, particularly ONT, may show elevated homopolymer rates due to systematic errors in homopolymer-region base calling. The module uses 5% and 10% warn and fail thresholds for the per-base fraction of bases in runs of 3+. These thresholds are intentionally set well below the expected genomic background of approximately 18–20%, meaning the module is designed to detect anomalously low homopolymer content (which would indicate data quality issues) rather than flagging normal genomic content. This is why the module is disabled by default for short-read data, where it would always trigger warnings due to normal genomic composition.

### **S2. Streaming Parallel Pipeline Architecture**

#### **S2.1 Pipeline Overview**

RastQC’s parallel pipeline is designed around a single-producer, multiple-consumer architecture implemented using Rust’s crossbeam library for lock-free bounded channels. The pipeline activates automatically for files exceeding 50 MB in size; smaller files are processed sequentially in a single thread to avoid the overhead of thread creation and state merging.

The pipeline consists of three phases. In Phase 1 (I/O), a dedicated reader thread decompresses and parses the input file, producing batches of sequences that are pushed into a bounded channel with a capacity of 2 × N, where N is the number of worker threads. The bounded channel provides natural backpressure: if all workers are busy, the reader blocks until a slot becomes available, preventing unbounded memory growth from read-ahead buffering. In Phase 2 (Processing), N worker threads each maintain a complete, independent set of all active QC module instances. Each worker pulls a batch from the channel, processes all reads in the batch through its local module instances, and requests the next batch. In Phase 3 (Merging), after the reader signals end-of-file and all workers have drained the channel, the main thread collects all worker module states and merges them pairwise using the merge_from() trait method. The merge is associative and commutative for all modules, ensuring that the final result is independent of the order in which workers are merged.

#### **S2.2 Adaptive Batch Sizing**

A fixed batch size (e.g., 16,384 reads) creates a severe memory imbalance between short-read and long-read processing. For 150 bp Illumina reads, 16K reads per batch consume approximately 2.4 MB of sequence data. For 20 kb PacBio reads, the same 16K reads consume approximately 320 MB per batch—which, multiplied by the channel capacity plus active workers, can require several gigabytes of buffer memory alone.

RastQC addresses this with adaptive batch sizing. During the first 1,000 reads, the mean read length is estimated. The target batch size in bytes is set to approximately 4 MB, and the number of reads per batch is computed as 4,000,000 divided by the estimated mean read length. The batch size is clamped to a minimum of 64 reads and a maximum of 65,536 reads. This produces the following approximate batch sizes for representative data types: for 76 bp Illumina reads, approximately 52,632 reads per batch; for 150 bp Illumina reads, approximately 26,667 reads per batch; for 5 kb ONT reads, approximately 800 reads per batch; and for 20 kb PacBio reads, approximately 200 reads per batch. This adaptive strategy keeps per-batch memory at approximately 4 MB regardless of read length, preventing the memory blowup observed with fixed batch sizes.

**Table S1.** Adaptive batch sizing: computed reads per batch for different read lengths, targeting ~4 MB per batch.

| **Mean Read Length** | **Reads per Batch** | **Approx. Batch Size** | **Typical Platform** |
| --- | --- | --- | --- |
| 76 bp | ~52,632 | ~4.0 MB | Illumina WES (short) |
| 126 bp | ~31,746 | ~4.0 MB | Illumina WES (medium) |
| 150 bp | ~26,667 | ~4.0 MB | Illumina WGS |
| 5,347 bp | ~748 | ~4.0 MB | ONT MinION |
| 18,814 bp | ~213 | ~4.0 MB | PacBio Revio HiFi |
| 50,000 bp | ~80 | ~4.0 MB | ONT ultra-long |
| 100,000 bp | ~64 (clamped) | ~6.4 MB | ONT ultra-long (max) |

#### **S2.3 Memory Model**

Total memory consumption during parallel processing can be decomposed into the following components. The baseline memory (M_base) covers the binary, I/O buffers, and overhead, and is approximately 15–20 MB. The per-worker module state (M_worker) covers all QC module accumulators per thread, including per-position arrays (1,000 positions × 94 quality values × 8 bytes = ~750 KB), duplication hash map (100K entries × ~80 bytes = ~8 MB), overrepresented sequence hash map (shared with duplication), and other module counters (~1 MB). Per worker, the total is approximately 10 MB for short-read mode and 11 MB for long-read mode. The channel buffer memory (M_channel) covers 2N batches of approximately 4 MB each, totaling 2 × 4 × 4 = 32 MB for 4 threads. The decompression buffer (M_decomp) covers the gzip/bzip2 decompression working memory at approximately 10–50 MB.

Total estimated memory = M_base + N × M_worker + M_channel + M_decomp. For 4 threads on short-read data: 20 + 4 × 10 + 32 + 30 = ~122 MB. The observed 49–50 MB for small sequential files (no parallel overhead) and 315–332 MB for large parallel files reflects this model, with additional memory attributed to hash map load factors, OS-level allocator overhead, and the input file’s memory-mapped I/O pages.

#### **S2.4 Thread Synchronization**

The pipeline uses minimal synchronization primitives to maximize throughput. The crossbeam bounded channel provides lock-free enqueue/dequeue operations in the common (non-full/non-empty) case. No mutexes are used during the processing phase—workers operate on entirely independent module instances. The only synchronization point is the merge phase after all workers complete, which is implemented as a sequential fold over worker states. Merge cost is O(M) per pair, where M is the total module state size (dominated by the duplication hash map). For 4 workers, the merge phase contributes less than 50 ms on typical datasets, which is negligible compared to the seconds-scale processing time.

### **S3. Extended Benchmark Data**

#### **S3.1 Complete Timing Breakdown**

Table S2 provides the full timing data for all benchmarked files, including user time and system time alongside wall-clock time. User time reflects total CPU time across all threads, while system time reflects kernel-level I/O operations. The ratio of user time to wall-clock time provides a measure of parallelization efficiency; a ratio close to N (the number of threads) indicates near-perfect parallel scaling.

**Table S2.** Complete timing breakdown for all benchmarked files (4 threads).

| **Tool** | **File** | **Size (MB)** | **Reads** | **Real (s)** | **User (s)** | **Sys (s)** | **RSS (MB)** | **Speedup** |
| --- | --- | --- | --- | --- | --- | --- | --- | --- |
| FastQC | DRR609229_1 | 22 | 720K | 3.50 | 3.45 | 3.45 | 425 | — |
| RastQC | DRR609229_1 | 22 | 720K | 1.99 | 1.93 | 1.93 | 49 | **1.8×** |
| FastQC | DRR609229_2 | 23 | 720K | 3.51 | 3.44 | 3.44 | 424 | — |
| RastQC | DRR609229_2 | 23 | 720K | 2.04 | 2.00 | 2.00 | 50 | **1.7×** |
| FastQC | ERR5897746_1 | 320 | 4.3M | 15.56 | 15.49 | 15.49 | 446 | — |
| RastQC | ERR5897746_1 | 320 | 4.3M | 4.81 | 4.76 | 4.76 | 332 | **3.2×** |
| FastQC | ERR5897746_2 | 327 | 4.3M | 15.57 | 15.49 | 15.49 | 442 | — |
| RastQC | ERR5897746_2 | 327 | 4.3M | 4.81 | 4.76 | 4.76 | 330 | **3.2×** |
| FastQC | DRR013000_1 | 1,430 | 24.8M | 51.75 | 51.69 | 51.69 | 434 | — |
| RastQC | DRR013000_1 | 1,430 | 24.8M | 19.62 | 19.57 | 19.57 | 315 | **2.6×** |
| FastQC | ALL_SHORT | — | — | 55.74 | 55.69 | 55.69 | 1,068 | — |
| RastQC | ALL_SHORT | — | — | 22.25 | 22.20 | 22.20 | 825 | **2.5×** |
| FastQC | ONT E. coli | 406 | 76K | 14.56 | 14.51 | 14.51 | 854 | — |
| RastQC | ONT E. coli | 406 | 76K | 3.12 | 3.07 | 3.07 | 1,257 | **4.7×** |
| RastQC-LR | ONT E. coli | 406 | 76K | 3.13 | 3.08 | 3.08 | 1,275 | **4.7×** |
| FastQC | PacBio E. coli | 281 | 42K | 17.57 | 17.51 | 17.51 | 702 | — |
| RastQC | PacBio E. coli | 281 | 42K | 2.72 | 2.67 | 2.67 | 670 | **6.5×** |
| RastQC-LR | PacBio E. coli | 281 | 42K | 2.71 | 2.67 | 2.67 | 676 | **6.5×** |

#### **S3.2 Throughput Analysis**

Table S3 presents throughput metrics computed from the benchmark data, expressed in megabytes per second and millions of reads per second. These metrics normalize for file size and read count, providing a more direct comparison of processing efficiency.

**Table S3.** Throughput comparison between RastQC and FastQC.

| **File** | **FastQC MB/s** | **RastQC MB/s** | **FastQC Mreads/s** | **RastQC Mreads/s** | **MB/s Ratio** | **Mreads/s Ratio** |
| --- | --- | --- | --- | --- | --- | --- |
| DRR609229 R1 (22 MB) | 6.3 | 11.1 | 0.206 | 0.362 | **1.8×** | **1.8×** |
| DRR609229 R2 (23 MB) | 6.6 | 11.3 | 0.205 | 0.353 | **1.7×** | **1.7×** |
| ERR5897746 R1 (320 MB) | 20.6 | 66.5 | 0.273 | 0.884 | **3.2×** | **3.2×** |
| ERR5897746 R2 (327 MB) | 21.0 | 68.0 | 0.273 | 0.884 | **3.2×** | **3.2×** |
| DRR013000 R1 (1,430 MB) | 27.6 | 72.9 | 0.479 | 1.263 | **2.6×** | **2.6×** |
| ONT E. coli (406 MB) | 27.9 | 130.1 | 0.005 | 0.024 | **4.7×** | **4.7×** |
| PacBio E. coli (281 MB) | 16.0 | 103.3 | 0.002 | 0.015 | **6.5×** | **6.5×** |

Several trends emerge from the throughput analysis. FastQC’s throughput in MB/s increases with file size (from 6.3 MB/s on a 22 MB file to 27.9 MB/s on a 406 MB file), reflecting the diminishing impact of JVM startup overhead as a fraction of total runtime. RastQC’s throughput also increases with file size but reaches much higher absolute values (up to 130.1 MB/s on ONT data), reflecting the efficiency of the streaming parallel pipeline. The throughput ratio is relatively constant in the reads-per-second metric (matching the speedup ratios), confirming that the speedup is consistent across different read lengths and file sizes.

#### **S3.3 Long-Read Module Overhead Analysis**

A key design question for RastQC was whether the three additional long-read modules would impose meaningful overhead. Table S4 compares RastQC performance with and without the --long-read flag to quantify the exact overhead of the additional modules.

**Table S4.** Overhead of --long-read flag (3 additional modules) on long-read datasets.

| **File** | **RastQC (12 modules)** | **RastQC --long-read (15 modules)** | **Δ Time** | **Δ Time %** | **Δ RSS** |
| --- | --- | --- | --- | --- | --- |
| ONT E. coli (406 MB) | 3.12 s | 3.13 s | +0.01 s | +0.3% | +18 MB (+1.4%) |
| PacBio E. coli (281 MB) | 2.72 s | 2.71 s | −0.01 s | −0.4% | +6 MB (+0.9%) |

The results demonstrate that the long-read modules add negligible overhead: less than 0.4% in wall-clock time and less than 1.4% in memory. On PacBio data, the --long-read variant was actually 0.01 seconds faster than the base variant, which is within measurement noise. The memory increase of 6–18 MB is attributable to the homopolymer run-length tracking matrix (4 bases × 20 run lengths × 8 bytes per counter × N workers) and the quality-stratified length histograms. This minimal overhead validates the design decision to integrate long-read modules directly into the streaming pipeline rather than implementing them as a separate processing pass.

### **S4. Output Concordance Methodology and Detailed Results**

#### **S4.1 Concordance Test Design**

To validate RastQC’s correctness, we performed a systematic comparison of module-level PASS/WARN/FAIL calls between RastQC v0.1.0 and FastQC v0.12.1 on a diverse set of datasets spanning five model organisms. For each dataset, both tools were run with default settings, and the resulting summary.txt files were compared. The comparison was restricted to the 11 modules shared between both tools (FastQC does not implement modules 12–15, and RastQC’s K-mer Content module is module 12, corresponding to FastQC’s module 11).

#### **S4.2 Test Datasets**

The concordance test used 10 datasets: the 5 short-read benchmark files described in the main text (DRR609229 R1, DRR609229 R2, ERR5897746 R1, ERR5897746 R2, DRR013000 R1) plus 5 additional model organism datasets from the European Nucleotide Archive. The model organism datasets were selected to cover a broad range of GC content, sequence complexity, and read characteristics, ensuring that concordance was not specific to a narrow class of inputs.

**Table S5.** Datasets used for concordance testing.

| **Organism** | **Strain/Assembly** | **GC Content** | **Genome Size** | **Notes** |
| --- | --- | --- | --- | --- |
| E. coli | *Escherichia coli K-12* | ~50.8% | ~4.6 Mb | Prokaryote, moderate GC |
| S. cerevisiae | *Saccharomyces cerevisiae S288C* | ~38.3% | ~12.1 Mb | Eukaryote, low GC |
| D. melanogaster | *Drosophila melanogaster* | ~42.5% | ~143 Mb | Complex eukaryote, repetitive |
| M. musculus | *Mus musculus C57BL/6J* | ~42.4% | ~2.7 Gb | Mammalian, large genome |
| H. sapiens | *Homo sapiens GRCh38* | ~40.9% | ~3.1 Gb | Mammalian, reference |

#### **S4.3 Concordance Results**

Across all 10 datasets and 11 shared modules, a total of 55 module-level PASS/WARN/FAIL comparisons were performed (the Per Tile Sequence Quality module is excluded from comparison for datasets lacking tile information, but all 5 short-read benchmark files contained tile information). All 55 comparisons produced identical results, yielding 100% concordance.

This concordance was achieved through careful reverse-engineering of FastQC’s internal algorithms, particularly in three modules where subtle implementation details significantly affect the output.

##### **S4.3.1 Per Sequence GC Content Concordance**

The GC Content module required the most careful implementation to achieve concordance. Three implementation details were critical. First, FastQC uses a GCModel class that distributes fractional counts across bins when a read’s GC percentage falls between integer bins. For example, a read with 45.3% GC contributes 0.7 counts to bin 45 and 0.3 counts to bin 46. RastQC implements this exact fractional binning. Second, the theoretical normal curve is fitted using the sum of the GC distribution array (which, due to fractional binning, may not equal the raw sequence count) as the denominator for standard deviation computation. Using the raw sequence count instead produces slightly different standard deviations, which can change the PASS/WARN/FAIL status at the boundary. Third, the mean GC is estimated using bidirectional mode averaging rather than the arithmetic mean of the distribution, which improves robustness for bimodal or skewed GC distributions.

##### **S4.3.2 Sequence Duplication Levels Concordance**

The Duplication module’s iterative binomial correction is FastQC’s most mathematically complex algorithm. The correction adjusts observed duplication counts to estimate the true duplication distribution that would be observed if all sequences (not just the first 100K unique) had been tracked. The formula iteratively solves: expected_observed[d] = total_unique × C(T/total_unique, d) × p^d × (1-p)^(T/total_unique - d), where T is total sequences, p is the probability of observing a given unique sequence, and d is the duplication level. RastQC implements the exact same iterative solver with the same convergence criteria and edge-case handling to ensure concordance.

##### **S4.3.3 Overrepresented Sequences Concordance**

The Overrepresented Sequences module required matching FastQC’s exact denominator for percentage calculations. FastQC uses the total number of sequences in the file (not the count of sequences processed before the hash map was frozen) as the denominator. This seems intuitive but differs from the behavior one might expect if the hash map is frozen early: a sequence observed 500 times out of 100K tracked sequences (0.5% of tracked) might constitute only 0.05% of a 1M-read file. Using the total count as the denominator produces the correct estimate of the sequence’s true prevalence. RastQC matches this behavior exactly.

### **S5. Compilation, Deployment, and Configuration Details**

#### **S5.1 Rust Compilation Profile**

RastQC is compiled using Rust’s release profile with additional optimizations enabled in the Cargo.toml configuration. Link-time optimization (LTO) is set to “fat” mode, which performs whole-program optimization across all crates, producing a smaller and faster binary at the cost of longer compilation times. The optimization level is set to 3 (opt-level = 3), which enables aggressive inlining, loop unrolling, and vectorization. Code generation units are set to 1, which enables maximum cross-module optimization. The panic strategy is set to “abort” rather than “unwind,” which eliminates unwinding tables and reduces binary size by approximately 10–15%. With these settings, the resulting static binary is 2.1 MB on the ARM64 target used for benchmarking.

#### **S5.2 Dependency Tree**

RastQC’s core functionality depends on the following Rust crates. The noodles crate (v0.85+) provides BAM/SAM format parsing. The flate2 crate provides gzip decompression using the miniz_oxide pure-Rust backend, which avoids a system library dependency on zlib. The bzip2 crate provides bzip2 decompression. The crossbeam crate provides lock-free bounded channels for the parallel pipeline. The clap crate provides command-line argument parsing. The serde and serde_json crates provide JSON serialization for MultiQC JSON output. Optional dependencies, gated behind the nanopore feature flag, include the hdf5 crate for Fast5 format support and the arrow crate for POD5 (Apache Arrow IPC) format support; these require the HDF5 system library when enabled.

#### **S5.3 Container Deployment**

RastQC’s static binary design makes it particularly well-suited for containerized deployment. A minimal Docker image can be constructed from the scratch or alpine base images with only the RastQC binary copied in, producing images as small as 2.5 MB (scratch + binary) or approximately 8 MB (alpine + binary). By contrast, a FastQC container requires a Java runtime environment (typically 200+ MB), resulting in images of 250+ MB. This 100× reduction in image size significantly accelerates container pull times in cloud-based workflow systems such as AWS Batch, Google Cloud Life Sciences, and Terra, where hundreds of container pulls may occur per pipeline run. A sample Dockerfile is provided in the repository.

#### **S5.4 Command-Line Interface**

RastQC provides a comprehensive command-line interface designed for both interactive use and automated pipeline integration. The core options mirror FastQC’s interface for drop-in compatibility. Table S6 summarizes the key command-line options and their descriptions.

**Table S6.** Key RastQC command-line options.

| **Option** | **Description** |
| --- | --- |
| -o, --outdir <DIR> | Output directory for results (default: current directory) |
| -t, --threads <N> | Number of processing threads (default: number of logical CPUs) |
| --long-read | Enable 3 additional long-read QC modules (N50, Quality Stratified Length, Homopolymer Content) |
| --multiqc-json | Generate native MultiQC JSON output alongside standard format |
| --serve [PORT] | Start built-in web server for interactive report viewing (default port: 8080) |
| --exit-code | Exit with non-zero status if any module reports FAIL (for pipeline gating) |
| --time | Print per-step timing breakdown (QC, report generation, I/O write) |
| --adapters <FILE> | Custom adapter sequence file (overrides built-in 6-adapter set) |
| --contaminants <FILE> | Custom contaminant sequence file (overrides built-in 15-entry set) |
| --limits <FILE> | Custom pass/warn/fail threshold file (overrides all module thresholds) |
| --nogroup | Disable adaptive base grouping (report every position individually) |
| --min-length <N> | Minimum read length to process (shorter reads are skipped) |
| --format <FMT> | Force input format (fastq, bam, sam, fast5, pod5; default: auto-detect) |

### **S6. Rust Language Design Considerations**

The choice of Rust as the implementation language for RastQC was driven by several technical factors that directly impact performance, correctness, and deployment characteristics in the bioinformatics context.

#### **S6.1 Zero-Cost Abstractions**

Rust’s trait system enables polymorphism (e.g., the QCModule trait shared by all 15 modules) without runtime virtual dispatch overhead. When the compiler can determine the concrete type at compile time, trait method calls are monomorphized—inlined as direct function calls with zero overhead compared to manually duplicated code. This allows RastQC to maintain a clean, extensible module architecture without paying the performance penalty typically associated with object-oriented designs in languages like Java (where interface dispatch involves vtable lookups and prevents inlining).

#### **S6.2 Memory Safety Without Garbage Collection**

Rust’s ownership and borrowing system guarantees memory safety at compile time without requiring a garbage collector. This has two performance implications for RastQC. First, there are no garbage collection pauses during processing, which eliminates the unpredictable latency spikes that Java applications experience (FastQC’s JVM may trigger GC pauses of 10–100 ms during processing). Second, memory is freed deterministically when values go out of scope, which means that per-batch sequence data is freed immediately after processing rather than accumulating until the next GC cycle. This contributes to RastQC’s lower memory usage, particularly on small files where the JVM’s generational garbage collector retains substantial amounts of dead object memory between collections.

#### **S6.3 Fearless Concurrency**

Rust’s type system enforces thread safety at compile time through the Send and Sync marker traits. A type is Send if it can be safely transferred to another thread; it is Sync if it can be safely shared between threads via references. The compiler rejects any program that attempts to share mutable state between threads without proper synchronization. This guarantee allowed RastQC’s parallel pipeline to be implemented with confidence that no data races exist—a guarantee that would require extensive runtime testing and careful code review in C++ or Java. In practice, this meant that the parallel pipeline was correct on the first implementation attempt, without the subtle concurrency bugs that plague multi-threaded bioinformatics tools.

#### **S6.4 Static Linking and Cross-Compilation**

Rust’s compiler toolchain supports static linking of all dependencies (including the Rust standard library) into a single binary with no external shared library dependencies. This produces the 2.1 MB static binary that characterizes RastQC’s deployment story. Cross-compilation is supported via the rustup target system: compiling for Linux x86_64 from a macOS ARM64 host requires only adding the target triple and specifying it at build time. This enables building release binaries for all major platforms from a single CI/CD pipeline, without requiring native build machines for each target architecture.

### **S7. Test Suite Details**

RastQC includes a comprehensive test suite of 36 tests (25 unit tests and 11 integration tests) that validate correctness at both the module level and the end-to-end pipeline level.

#### **S7.1 Unit Tests**

Unit tests validate individual module logic in isolation. Each QC module has dedicated tests that process synthetic or known-content sequences and verify that the module’s statistics, thresholds, and PASS/WARN/FAIL decisions match expected values. Specific unit test categories include I/O parser tests that verify correct parsing of FASTQ, compressed FASTQ, BAM, and SAM formats, including edge cases such as empty files, truncated records, and non-standard quality encodings. Module accumulator tests verify that per-position counters, histograms, and hash maps are updated correctly for known input sequences. Merge tests verify that the merge_from() operation produces identical results regardless of how the input is partitioned across workers. This is tested by processing the same input sequentially and in parallel with varying numbers of workers, asserting identical final module states. Threshold tests verify that PASS/WARN/FAIL decisions are correct at boundary values for each module.

#### **S7.2 Integration Tests**

Integration tests validate the complete pipeline from file input to report output. Each integration test runs RastQC on a known dataset and checks the output summary.txt against expected PASS/WARN/FAIL calls, verifies that the HTML report is valid and contains expected sections, verifies that the fastqc_data.txt file is parseable by a reference MultiQC implementation, and checks that ZIP archive structure matches the expected layout. Additional integration tests verify the multi-sample summary dashboard generation when processing multiple files simultaneously, the --multiqc-json output against the expected JSON schema, the --exit-code behavior (zero exit for all-PASS, non-zero exit for any FAIL), and the --serve web server startup and response to HTTP requests.

#### **S7.3 Continuous Integration**

The test suite is executed on every commit via GitHub Actions, running on Linux (Ubuntu 22.04, x86_64), macOS (14, ARM64), and Windows (Server 2022, x86_64) to ensure cross-platform correctness. The CI pipeline also runs the concordance comparison against FastQC v0.12.1 on a subset of benchmark files to detect any regressions in output compatibility.

### **S8. Output Format Specifications**

#### **S8.1 Directory Structure**

For each input file (e.g., sample.fastq.gz), RastQC produces the following output directory structure within the specified output directory. The main output directory contains sample_fastqc.html (the self-contained HTML report), sample_fastqc.zip (the ZIP archive for MultiQC compatibility), and if --multiqc-json is specified, sample_fastqc_multiqc.json. Inside the ZIP archive, the structure is sample_fastqc/fastqc_data.txt (tab-separated module data), sample_fastqc/summary.txt (per-module PASS/WARN/FAIL status), and sample_fastqc/fastqc_report.html (duplicate of the HTML report). When multiple files are processed, a summary directory is additionally created containing multisample_summary.html (interactive multi-sample dashboard) and multisample_summary.tsv (tab-separated matrix of module statuses across all samples).

#### **S8.2 fastqc_data.txt Format**

The fastqc_data.txt file uses a section-based format where each module’s data is enclosed between a header line (>>Module Name\tpass/warn/fail) and a footer line (>>END_MODULE). Within each section, data is tab-separated with a header row prefixed by #. Column names, data types, and value ranges exactly match FastQC’s output to ensure that MultiQC’s FastQC parser can process RastQC output without modification. The Basic Statistics section includes the Filename, File type, Encoding, Total Sequences, Total Bases, Sequences flagged as poor quality, Sequence length, and %GC fields.

#### **S8.3 MultiQC JSON Format**

The --multiqc-json flag produces a structured JSON file designed for direct consumption by MultiQC’s JSON input module. The JSON structure includes a top-level object with report_metadata (tool name, version, timestamp, command), sample_metadata (per-sample file information), and module_data (per-module results keyed by module name). Each module’s data includes the status (pass/warn/fail), raw data arrays or objects, and any module-specific metadata. This format eliminates the text-parsing overhead required by MultiQC’s FastQC parser when processing the tab-separated fastqc_data.txt format, reducing MultiQC processing time for large projects with hundreds or thousands of samples.

#### **S8.4 HTML Report Structure**

RastQC’s HTML reports are fully self-contained, with all CSS, JavaScript, and SVG charts inlined. No external resources are loaded, ensuring that reports render correctly offline and in restricted network environments. Charts are rendered as inline SVG elements with complete axis labels, tick marks, and legends, providing publication-quality visualizations directly from the report. The report includes a sidebar navigation with module-level PASS/WARN/FAIL indicators (color-coded green, orange, and red), enabling rapid identification of problematic modules. The multi-sample summary dashboard uses an interactive table with sortable columns and color-coded cells for cross-sample comparison.
